# Affinity and Conformations of Ca^2+^-Calmodulin Bound to Peptides Containing the Calmodulin-Binding Motif of HIV-1 Matrix Protein

**DOI:** 10.64898/2026.10.19.760223

**Authors:** Arvin Saffarian Delkhosh, Adeyemi Ogunbowale, Elaheh Hadadianpour, Kazi Mustafa Mahmud, Peter P. Borbat, Elka R. Georgieva

## Abstract

**Background:** Protein–protein interactions enable viruses to adapt and replicate by modifying host factors. Thermodynamic characterization of viral–host complexes can reveal viral mechanisms, broader principles of protein interactions, and properties relevant to anti-HIV-1 inhibitor development. Here, we examine the binding of the HIV-1 matrix (MA) protein to host calmodulin (CaM). MA, the N-terminal domain of Gag (group-specific antigen), directs Gag to the plasma membrane for viral assembly and maturation and may recruit CaM to facilitate this trafficking. However, the MA–CaM interaction remains poorly characterized. MA contains a noncanonical N-terminal CaM-binding motif with hydrophobic regions that may mediate CaM binding and a central, positively charged KKK cluster.

**Objective:** We aimed to determine the binding affinities of MA to calcium-bound CaM (Ca^2+^-CaM), elucidate protein regions that are critical to the heterocomplex formation, and monitor the conformational alterations in Ca^2+^-CaM upon binding to MA.

**Methods:** We studied Ca^2+^-CaM binding to synthetic MA peptides representing full-length wild type (FL WT), a C-terminal truncation lacking the KKK cluster, and an FL K→N/K→H/K→Q triple mutant of the MA CaM-binding motif. The peptides were C-terminally labeled with Cy3, and FL CaM S39C with Cy5. Binding affinity was measured by ensemble Förster resonance energy transfer (eFRET) using 100 nM Cy3-MA and increasing Cy5-Ca^2+^-CaM concentrations in the range of 50 nM to 2.5 µM. Conformational changes in doubly spin-labeled Ca^2+^-CaM upon MA binding were probed by double electron-electron resonance (DEER) spectroscopy.

**Results:** Cy3 quenching and FRET analyses of complexes with Cy5-Ca^2+^-CaM yielded dissociation constants of ∼80 nM for FL WT MA, ∼65 nM for the truncation, and ∼250 nM for the triple mutant, with corresponding binding free energies of ∼−9.8 to ∼−9 kcal/mol. These values indicate stable, specific complexes and show that the KKK cluster contributes little to MA–Ca^2+^-CaM binding. DEER measurements of Ca^2+^-CaM labeled at S39C and A103C showed that FL WT MA binding shifted the ∼5.3 nm interspin distance of extended Ca^2+^-CaM to a broader ∼1.8–3 nm distribution, indicating a major CaM restructuring.

**Conclusion:** These findings define key energetic and structural features of MA-Ca^2+^-CaM interactions and clarify how HIV-1 proteins exploit CaM.

## 1. Introduction

Protein-protein interactions are essential for virus adaptation and replication within host cells. Viral proteins bind host factors to take over functions, change their location, or target them for degradation^1, 2^. Elucidating the formation and structure of viral-host protein complexes deepens our understanding of the biochemical processes underlying viral mechanisms and general protein interactions.

Among many other interactions, of importance are those of the HIV-1 proteins with host calmodulin (CaM). Amino acid (aa) sequence analysis indicates that most HIV-1 proteins possess CaM-binding motifs, although they are non-canonical and differ significantly from those in host proteins^3–7^. Direct association of HIV-1 Matrix (MA), Nef, Tat, etc. proteins with CaM have been reported, and it is believed that these interactions aid HIV-1 proteins’ trafficking, insertion in the membrane, regulating cellular apoptosis and contribute to the CD4+-lymphocytes deterioration^8–14^. Notably, it was reported that, compared to non-infected cells, the HIV-1 infected cells have higher CaM levels^15^, pointing to the significant role of CaM in the virus life cycle. Recently, our lab provided the first experimental evidence of a complex formation between the soluble form (which we discovered earlier^16, 17^) of the HIV-1 accessory protein Vpu and calcium-bound CaM (Ca^2+^-CaM); we estimated that the free binding energy (ΔG) of Vpu-Ca^2+^-CaM interactions is near or slightly less than – 10 kcal/mol, suggesting a stable complex formation. Based on these findings, we proposed that this interaction could facilitate the trafficking of Vpu in the cell^18, 19^. CaM is a Ca^2+^-sensor protein, ubiquitously expressed at high levels and participating in control of signaling and metabolic processes^20–22^. HIV-1 appears to have evolved efficient mechanisms to exploit CaM. Despite their importance in regulating cellular physiology, interactions between HIV-1 proteins and CaM remain insufficiently studied.

Our focus here is on the mechanisms of HIV-1 MA protein-CaM complex formation. MA is the N-terminal domain of the Gag (group-specific antigen) polypeptide containing about 132 aa-s, which folds in a globular structure ^23, 24^ (Figure 1A, B). Through MA, the Gag protein is targeted and anchored to the plasma membrane (PM)^25, 26^. The MA domain interacts with PM via electrostatic interactions involving the highly basic aa-s region (HBR), residues 18–33 (Figure 1B)^27^. The Gag polyprotein is synthesized in the cytoplasm and plays critical functions in HIV-1 replication, assembly, budding, and release of viral particles^28, 29^. Because CaM and Gag were found to co-localize in the cytosol, it was suggested that the HIV-1 Gag protein recruits CaM for trafficking toward the PM to the site of virus assembly^11^. The Gag-CaM association occurs via the MA domain and is Ca^2+^-dependent^10^. Structural studies revealed that conformational changes in both the MA and CaM underly their complex formation with helix 1 of MA (Figure 1C) adopting an open conformation^11^. The stretch of residues 8–43, including the membrane-binding HBR, in MA was identified as the entire CaM-binding region (Figure 1), although peptides containing residues 11-28 and 31-46 also bind CaM but with much reduced affinity^30^.

**Figure 1.**
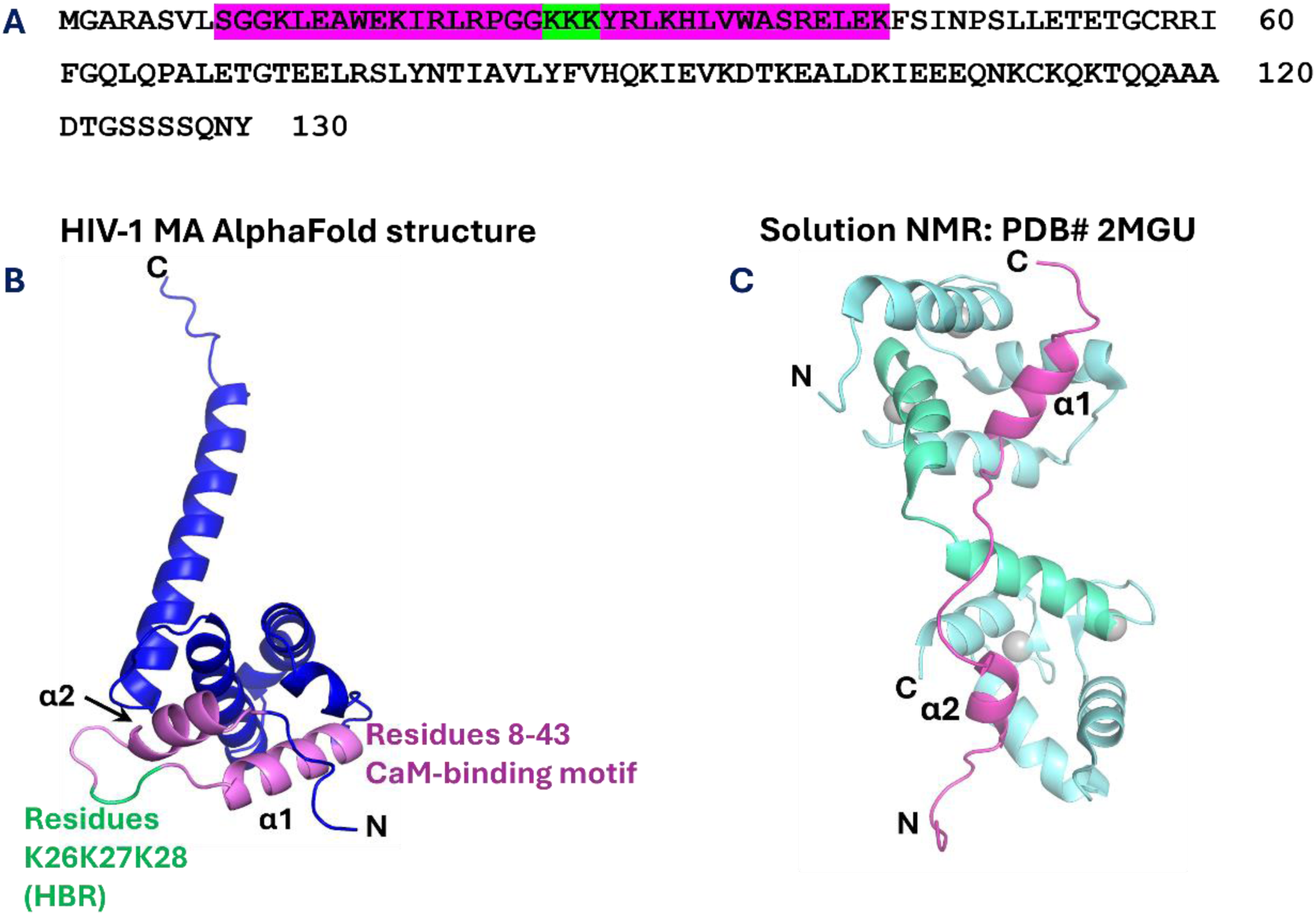
Structural features of the HIV-1 MA protein and its complex with Ca^2+^-CaM. (A) Amino acid sequence of the FL MA construct. The CaM-binding motif (residues 8–43) and the K26–K28 lysine cluster are highlighted in magenta and green, respectively. (B) AlphaFold-predicted structure of the MA construct, with the CaM-binding motif shown in magenta and residues K26– K28 shown in green. The CaM-binding motif folds into two helices (α1 and α2) that are arranged in antiparallel manner. (C) Solution NMR structure of the MA CaM-binding motif (magenta) in complex with Ca^2+^-CaM (cyan cartoon and Ca^2+^ in gray spheres). The central CaM helix, which adopts a kinked conformation upon MA binding, is shown in cyan-green. The two short helices, α1 and α2, of the MA CaM-binding motif form more extended conformation and contact the CaM lobes. Ca^2+^-CaM and MA bind in an antiparallel orientation: the N-terminal α1 helix binds the C-terminal CaM lobe, whereas α2 binds the N-terminal lobe.

In our studies, we tested the affinity of Ca^2+^-CaM to three peptides (Figure 2, Table 1) that are (1) wild type (WT) peptide corresponding to the FL CaM-binding motif of the MA protein (residues 8-43 in FL MA protein—all residue numbering throughout the paper is in FL MA) containing a triplet of Lys residues (K26K27K28) of the HBR, (2) the same peptide but carrying a triple mutation of the HBR region (K26N/K27H/K28Q) aa numbering in FL MA protein)—referred as MA-Mut, and (3) a peptide corresponding to the C-terminal region of MA’s CaM-binding motif (residues 29-43, aa numbering in FL MA) that does not have the KKK positive charges—referred as MA-Ct. We aimed to understand the role of CaM-binding motif regions, including the hydrophobic residues in the N-terminal and C-terminal part, as well as the clustered positive residues in HBR on MA-Ca^2+^-CaM interactions. The latter has not been evaluated to date; however, it is important to know whether these residues are essential to MA-CaM binding by adding electrostatic interactions or whether the MA-Ca^2+^-CaM complex is stabilized predominantly by hydrophobic interactions.

**Figure 2.**
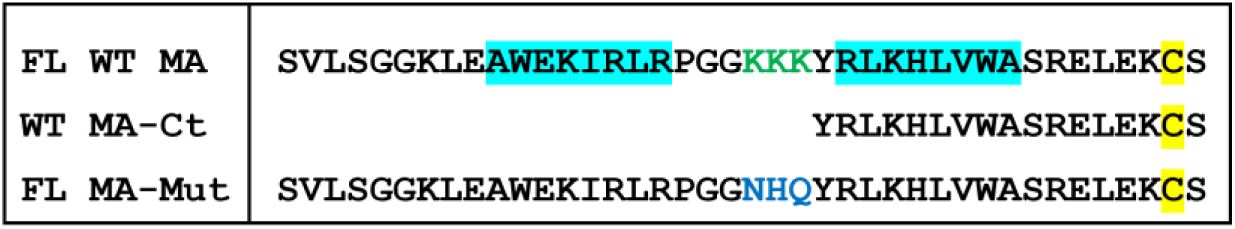
The amino acid sequences of the peptides containing the MA CaM-binding domain used in this study. The imperfect repeat regions in the FL WT MA peptide are highlighted in cyan and the KKK cluster in the HBR are colored in green.

**Table 1.** MA peptide and CaM constructs produced and used in this study.

| Construct | Number of residues in the construct | Mutation(s) | Tag | MW (kDa) | $\epsilon_{280}$ ($M^{-1} \text{ cm}^{-1}$) | Label |
| --- | --- | --- | --- | --- | --- | --- |
| FL WT MA | 40 | — | Untagged | 4,638.50 | 12,490 | Cy3 |
| WT MA-Ct | 17 | — | Untagged | 2,118.48 | 6,990 | Cy3 |
| FL MA-Mut | 40 | K26N/K27H/K28Q | Untagged | 4,633.36 | 12,490 | Cy3 |
| CaM S39C | 171 | S39C | N-terminal His<sub>10</sub> | 19,402.40 | 2,980 | Cy5 |
| CaM S39C/A103C | 171 | S39C/A103C | N-terminal His<sub>10</sub> | 19,434.46 | 2,980 | MTSL |

To characterize the MA peptides-Ca^2+^-CaM complexes formation, we used ensemble FRET (eFRET) on Cy3 (donor)-labeled MA peptides and Cy5 (acceptor)-labeled CaM at engineered single cysteine residues, similarly to the Vpu-Ca^2+^-CaM study^19^. For all studied MA peptides, we obtained *K*_d_‘s in the range of 65 nM-80 nM, and somewhat greater, i.e., 250 nM for the MA-Mut, corresponding to binding energies of ∼ - 9.8-9 kcal/mol. Strikingly, these binding parameters are close to those, which we obtained for the Vpu-Ca^2+^-CaM complex, suggesting similar binding affinity, although based on dissimilar mechanisms, of the two HIV-1 proteins (Vpu and MA) to Ca^2+^-CaM. Our results strongly suggest that the removal of the positive KKK residues in MA has no or only a minor effect on the MA-Ca^2+^-CaM complex formation, as the binding affinity, characterized by the estimated dissociation constant *K*_d_ and free binding energy (*ΔG*) were almost unaffected by MA CaM-binding motif truncation or mutations.

We applied DEER spectroscopy to spin-labeled at residues S39C/A103C Ca^2+^-CaM^18^ and detected the transition from open to deemed closed Ca^2+^-CaM conformations upon binding to the FL MA peptide, which is in general agreement with NMR and our prior results for the Vpu-Ca^2+^-CaM^12,18^.

This study expands our understanding of the nature of HIV-1 proteins complexes with Ca^2+^-CaM, which play physiological roles in the infected cells. These findings also broaden our understanding of viral protein complexes with Ca^2+^-CaM and how viruses exploit Ca^2+^-binding proteins to support their functions.

## 2. Materials and Methods

### 2.1. Protein designs, cloning, mutagenesis, expression and purification

This study used three HIV-1 matrix protein (MA) constructs: full-length wild-type MA (FL WT MA), the C-terminal region of MA spanning residues 29–43 (MA-Ct), and a full-length MA mutant carrying the K21N/K22H/K23Q substitutions (FL MA-mut). These peptides were commercially synthesized (GenScript, Inc.) with >95% purity and provided in lyophilized form. Upon receipt, we reconstituted each peptide in 20 mM Tris-HCl (pH 7.4), 150 mM NaCl, 5% (v/v) glycerol and 5 mM DTT (dithiothreitol), then aliquoted and stored at −80 °C until use.

The CaM protein was the same as that used in our previous study on Vpu-CaM^18^. The single-cysteine CaM S39C and double-cysteine CaM S39C/A103C variants were expressed and purified following our previously established protocol^18^, with the S39C variant used for fluorescence labeling and the S39C/A103C variant used for spin labeling and DEER measurements.

The molar extinction coefficients (ε) at 280 nm, calculated from the amino acid sequences using ExPASy ProtParam, were 12,490 M^−1^ cm^−1^, 6,990 M^−1^ cm^−1^, 12490 M^−1^ cm^−1^ and 2,980 M^−1^ cm^−1^ for FL WT MA, WT MA-Ct, FL MA-Mut and CaM, respectively. These ε’s and absorbance at 280 nm were used to calculate peptide/protein concentrations in the following steps.

### 2.2. Cy3 and Cy5 fluorescence labeling of MA and CaM

For fluorescence experiments, the three MA constructs (FL WT MA, WT MA-Ct, and FL MA-Mut) were labeled with Cy3 maleimide, whereas the single-cysteine CaM S39C variant was labeled with Cy5 maleimide. Prior to labeling, the MA proteins were dialyzed to remove DTT present in the initial protein buffer. Dialysis was performed using 0.5–1 kDa molecular-weight cutoff (MWCO) Spectra/Por pre-wetted regenerated cellulose (RC) dialysis tubing against buffer containing 20 mM Tris/HCl (pH 7.4), 150 mM NaCl, 1 mM CaCl_2_, 50 μM TCEP, and 5% (v/v) glycerol. The dialysis buffer was replaced four times at 3 h intervals at 4°C to ensure all the DTT has been successfully removed. Following dialysis, the protein concentrations were determined using a NanoDrop OneC spectrophotometer (Thermo Scientific). Each MA construct was labeled with Cy3 maleimide (Lumiprobe) at a protein-to-dye molar ratio of 1:5. The labeling reaction was conducted at 22 °C for 3.5 h with continuous agitation using a Roto-Therm Mini rotator (Benchmark Scientific), followed by overnight incubation at 4 °C with continuous agitation. On the following day, the excess unreacted Cy3 maleimide dye was removed through dialysis against the buffer 20 mM Tris/HCl (pH 7.4), 150 mM NaCl, 1 mM CaCl_2_, 50 μM TCEP, and 5% (v/v) glycerol and the dialysis protocol followed the exact procedure described above. Throughout the labeling and all subsequent handling, samples were wrapped in aluminum foil to protect the cyanine dyes from light. The Cy3-labeled MA constructs were collected, and the final concentrations were determined.

The single-cysteine CaM S39C variant was labeled with Cy5 maleimide (Lumiprobe) using the same protein-to-dye molar ratio and labeling conditions described above for the MA constructs. Excess unreacted Cy5 dye was removed in the same way. The resulting Cy3-labeled MA constructs and Cy5-labeled CaM S39C were used for subsequent eFRET measurements.

### 2.3. Determination of Cy3 and Cy5 labeling efficiency

After removal of unreacted dye, the labeling efficiencies of the Cy3-labeled MA constructs and the Cy5-labeled CaM S39C variant were determined using a NanoDrop OneC spectrophotometer. Dye and protein concentrations were determined using the Protein and Labels function. For Cy3 maleimide, the dye concentration was determined from the absorbance at 555 nm using an ε of 150,000 M^−1^ cm^−1^:

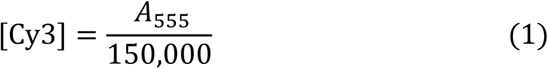

For Cy5 maleimide, the dye concentration was determined from the absorbance at 646 nm using an ε of 250,000 M^−1^ cm^−1^:

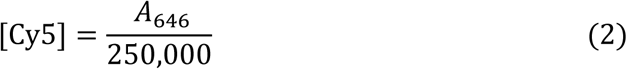

Because the fluorophores also contribute to the absorbance at 280 nm, protein concentrations were corrected using the manufacturer-provided correction factors (CF_280_) of 0.09 for Cy3 and 0.04 for Cy5. For the Cy3-labeled MA constructs, protein concentration was calculated as:

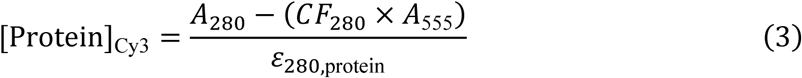

For Cy5-labeled CaM S39C, protein concentration was calculated as:

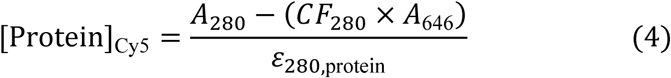

where ε_280,protein_ is the molar extinction coefficient of the respective protein at 280 nm (Table 1). The labeling efficiency was expressed as the molar ratio of bound dye to protein (degree of labeling, *D*/*P*):

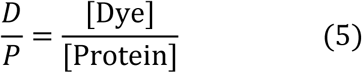

Where D is concentration of dye and P is concentration of protein.

The resulting labeling efficiencies were 0.54, 0.48, 0.35, and 0.55 for FL WT MA-Cy3, WT MA-Ct-Cy3, FL MA-Mut-Cy3, and CaM S39C-Cy5, respectively.

### 2.4. Ensemble FRET measurements of MA–Ca^2+^-CaM interactions

All eFRET experiments were performed in triplicate, with three independently prepared and measured samples, to assess the reproducibility and reliability of the results. The data presented in Figures 4, 5, 6, and 7 are expressed as the mean ± standard deviation (SD) calculated from these triplicate measurements. To ensure consistency across experiments, the mixing time between labeled MA variants and labeled Ca²⁺–CaM was carefully controlled, and all samples within each experimental series were analyzed under identical conditions. These precautions minimized potential experimental artifacts, including donor photobleaching and dilution effects, thereby reducing the likelihood of generating false eFRET signals.

All eFRET measurements were conducted at 25 °C using an FS5 spectrofluorometer (Edinburgh Instruments) equipped with a temperature-control system.

The following samples were prepared and eFRET experiments conducted on them:

a) To conduct quantitative analysis of MA variant-Ca^2+^-CaM binding, samples containing a mixture of 100 nM Cy3-labeled FL MA, MA-Ct, and FL MA-mut were mixed with increasing concentration of Cy5-labeled Ca^2+^-CaM at 0.05 µM, 0.1 µM, 0.2 µM, 0.35 µM, 0.45 µM, 0.6 µM, 0.85 µM, 1 µM, 1.5 µM, 2 µM, and 2.5 µM. The FRET data were collected using an FS5 spectrofluorometer (Edinburgh Instrument) with the donor excitation at 550 nm (3 nm excitation and emission bandwidth), and emission recorded from 555 to 800 nm. Each sample was prepared by mixing Cy3-MA variant with Cy5-Ca^2+^-CaM in the FRET buffer containing 1 mM CaCl_2_ in a cuvette. The protein mixtures were incubated at RT in a dark room for 10 minutes to allow protein interaction.
b) To evaluate the dynamic nature of the MA-Ca²⁺-CaM complexes, Cy3-FL MA was maintained at a constant concentration of 100 nM, whereas Cy5-labeled Ca²⁺-CaM was used at concentrations of 0.1 µM, 0.35 µM, 0.6 µM, 0.85 µM, 1.5 µM, and 2.5 µM. In selected samples, non-labeled Ca²⁺-CaM was subsequently added at final concentration of 10 µM. All samples were adjusted to a final volume of 1000 µL to maintain constant concentrations of the labeled proteins and non-labeled protein. Thus, the samples were prepared by initially combining 800 µL of FRET buffer (20 mM Tris pH 7.4, 150 mM NaCl, 1 mM CaCl_2_, 50 µM TCEP) with the appropriate volumes of Cy3-FL MA protein and Cy5-Ca²⁺-CaM, followed by a 10 min incubation period in a dark room at room temperature (RT) to allow complex formation. Non-labeled Ca²⁺-CaM was then added, and the buffer (20 mM Tris pH 7.4, 150 mM NaCl, 1 mM CaCl_2_, 50 µM TCEP) was used to bring the final volume to 1000 µL. The mixtures were incubated for an additional 5 min in dark at RT before eFRET measurements were acquired. Fluorescence spectra were collected using Cy3 donor excitation at 550 nm, with 1.5 nm excitation and emission bandwidths, and emission was recorded from 555 to 800 nm.

### 2.5. Determination of dissociation constants and data analysis

The dissociation constants (*K*_d_) for the MA-variant–Ca²⁺–CaM complexes were determined using Q-FRET quenching and KD-FRET analyses, following our previously established protocol^19^. For Q-FRET quenching, changes in donor (Cy3) fluorescence intensity upon incremental addition of acceptor (Cy5)-labeled Ca^2+^-CaM were analyzed using the nonlinear binding model described by Jiang et al^31^. Independently, *K*_d_ values were obtained through KD-FRET analysis, in which FRET efficiencies calculated according to Shresta et al^32^ were fitted to the quadratic binding equation reported by Yi et al^33^. Both approaches were applied to triplicate measurements, with the resulting *K*_d_ values expressed as the mean ± standard deviation. Data fitting and graphical analyses were conducted using OriginPro 2023 (OriginLab, Northampton, MA, USA). The quality of fit (R^2^) in all cases was greater than 92%.

### 2.6. Calmodulin spin labeling, DEER sample preparation, measurements, and data analysis

Ca^2+^-CaM with cysteine residues at positions S39C and A103C was tagged with the paramagnetic MTSL spin-label at protein-to-MTSL molar ratio of 1:15. The protein in 20 mM Tris/HCl pH 7.4, 150 mM NaCl, 1 mM CaCl_2_, and 5% glycerol buffer was mixed with MTSL dissolved in acetonitrile and incubated for 2 h at RT and then overnight at 4 °C under constant agitation. Next day, the unbound MTSL was removed by washing the protein with the same buffer in a concentrator with 10 kDa MWCO. The last washing buffer had the same composition but ≍80% D_2_O instead of H_2_O. Thereafter, a sample of spin-labeled Ca^2+^-CaM in ≍80% D_2_O buffer and 25% glycerol-*d*_8_ (w/v) at ≍40 µM final concentration was prepared. A second sample was made by mixing ca. 50 µM doubly spin-labeled Ca^2+^-CaM and 50 µM unlabeled FL WT MA peptide (after fully removing the DDT); then, the mixture was diluted 12 times using the 20 mM Tris/HCl pH 7.4, 150 mM NaCl, 1 mM CaCl_2_ in ≍80% D_2_O and incubated for about 10 min on ice. This dilution step was intended to promote the dissociation of FL WT MA peptide homooligomers at low total peptide concentration, thus making more MA monomers available to interact with the spin-labeled Ca^2+^-CaM. Next, the diluted sample was concentrated in a concentrator with 3 kDa MACO to the original volume before dilution and glycerol-*d*_8_ was added to 25% (w/v), so the final concentrations of the Ca^2+^-CaM and FL WT MA peptide were about 38 µM.

All double electron-electron resonance (DEER) measurements were conducted at temperature 60 K using a Bruker E580 EPR spectrometer equipped with a EN 5107D2 resonator operating in Q-band. Standard 4-pulse DEER setup^34^ was used with pumping performed at the low-field peak of the nitroxide spectrum. The raw data were reconstructed to distance distributions using Comparative Deer Analyzer 2.0 automated software application for reconstruction of distance distributions from the DEER data^35^.

### 2.7. Protein structure modeling

We used AlphaFold3^36^ to predict the structures of proteins and protein complexes. The predicted structures were subsequently visualized and analyzed using PyMOL.

## 3. Results

### 3.1. Ensemble FRET demonstrates the association of peptides derived from the MA CaM-binding motif with Ca^2+^-CaM

Our initial goal was by using eFRET to detect the MA peptides association with Ca^2+^-CaM. In these experiments we used 100 nM Cy3-labeled peptides that corresponded to the FL WT, C-terminal and FL mutant (K26N/K27H/K28Q) of MA CaM-binding motif (Figure 2, Table 1). The concentration was selected to maintain the MA peptides predominantly in their monomeric state, as we did for the Vpu protein^19^. The peptides in buffered solution were combined with Cy5-labeled Ca^2+^-CaM at concentrations ranging from 50 nM to 2.5 µM. eFRET arising from the formation of Cy3-MA peptide–Cy5-Ca^2+^-CaM complexes was monitored as a function of Cy5-Ca^2+^-CaM concentration (Figure 3). In each case, increasing the Cy5-Ca^2+^-CaM concentration produced a progressive decrease in Cy3 emission and a concomitant increase in Cy5 emission, consistent with Cy3-to-Cy5 energy transfer. These observations provide strong evidence that all MA peptides associate with Ca^2+^-CaM to form heterocomplexes. Fluorescence emission spectra of the Cy3-labeled MA peptides recorded in the absence of Cy5-Ca^2+^-CaM served as background signals and were subtracted from the signals of samples with increasing Cy5-Ca^2+^-CaM concentrations for further analysis.

**Figure 3.**
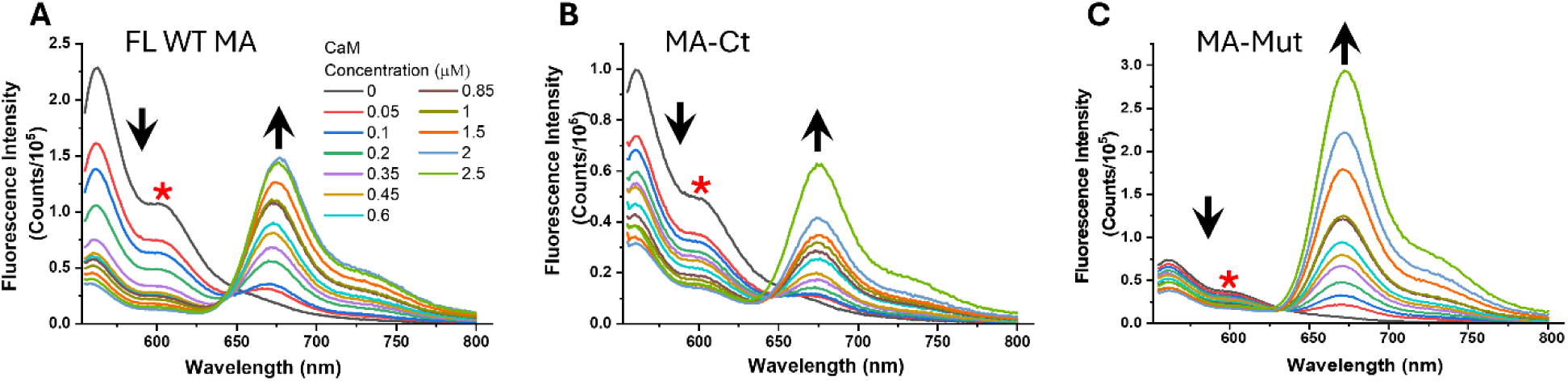
eFRET data for 100 nM Cy3-labeled MA peptides titrated with Cy5-labeled Ca^2+^-CaM over a concentration range of 0–2.5 µM. Spectra for FL WT MA, MA-Ct, and FL MA-Mut are presented in panels A, B, and C, respectively. Cy5-Ca^2+^-CaM concentrations are color-coded as indicated in panel A. Increasing Cy5-Ca^2+^-CaM concentrations produced progressive quenching of Cy3 emission between 550 and 612 nm, accompanied by enhanced Cy5 emission with a maximum near 675 nm, consistent with Cy3-to-Cy5 energy transfer. The decrease of Cy3 and increase of Cy5 emissions are designated with arrows in the corresponding directions, and the Cy3 emission wavelength (∼605 nm) which was used to estimate the *K*_d_‘s is designated with red asterisk.

### 3.2. Ensemble FRET data analysis yields dissociation constants (*K*_d_) of 65–250 nm and corresponding binding free energies of approximately −9.8 kcal/mol to −9.0 kcal/mol, consistent with the formation of specific, stable complexes between the MA peptides and Ca^2+^-CaM

Following our recent study on HIV-1 Vpu-Ca^2+^-CaM study^19^, we analyzed the eFRET data shown in Figure 3 adopting two methods based on Cy3 donor quenching upon Cy3-MA-Ca^2+^-CaM complexes formation that are:

(i) Quantitative FRET quenching^31^ (Q-FRET quenching) by fitting the data in Figure 3 for each of the MA peptide variants to the equation:

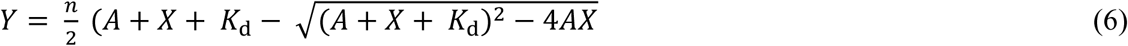

Where, *n* is a constant related to FRET efficiency (*E*_FRET_) between donor (MA peptide) and acceptor (CaM) – obtained from fittings, *A* is the constant concentration of donor (Cy3-MA peptide), and *X* is the varying concentration of acceptor (Cy5-Ca^2+^-CaM).

(ii) KD-FRET, previously developed to quantify protein–protein interactions in bacterial cells, relying on the *E*_FRET_ ^33^—For each series of samples containing 100 nM Cy3-MA variant and Cy5-Ca^2+^-CaM concentrations in the range of 100 nM to 2.5 µM, the values of *E*_FRET_ were calculated using equation (2)^37^:

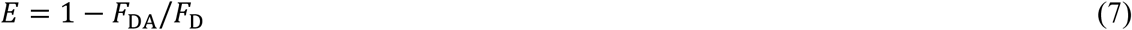

Where *E* is the FRET efficiency (*E*_FRET_), *F*_D_ is the fluorescence intensity of the Cy3 donor at about 610 nm without Cy5-Ca^2+^-CaM (in Figure 3, shown with asterisk) in the sample; and *F*_DA_ is the fluorescence intensity at the same peak maximum for each sample in the presence of Cy5-Ca^2+^-CaM.

Thereafter, the *K*_d_‘s were obtained by fitting these data to the equation:

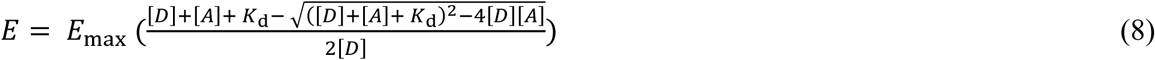

Where *E*_max_ is the maximum FRET efficiency (obtained from fittings), [*D*] is the concentration of donor-labeled protein (MA peptides), and [*A*] is the varying concentration of acceptor-labeled protein (Ca^2+^-CaM).

#### 3.2.1. The FL WT MA peptide binds to Ca^2+^-CaM with a K_d_ ∼80 nM corresponding to a binding energy of about −9.7 kcal/mol

The conducted binding analyses of Cy3-FL WT MA with Cy5-Ca^2+^-CaM yielded *K*_d_ values of 70.7 ± 9.8 nM by Q-FRET quenching and 91.6 ± 10.6 nM by KD-FRET (Figure 4) (Table 1). The calculated average *ΔG* was approximately −9.7 kcal/mol, consistent with formation of a stable protein-protein complex^38, 39^. These binding parameters indicate that the FL WT of the MA CaM-binding motif has relatively high affinity to Ca^2+^-CaM. The results are also consistent with earlier study, which found a *K*_d_ of about 25 nM for FL CaM-binding motif in MA protein^30^, yielding ΔG of ∼ −10.5 kcal/mol. Further, these findings suggest that the MA CaM binding motif-Ca^2+^-CaM complex is somewhat less stable compared to other CaM complexes with host proteins, which typically exhibit *ΔG* values ranging from −10 kcal/mol to −15 kcal/mol^40^. Notably, the binding parameters determined for the FL WT MA peptide closely match those recently obtained for the FL WT HIV-1 Vpu protein (∼ −9.8 kcal/mol)^19^. This similarity supports our earlier hypothesis that HIV-1 proteins may have evolved to exploit more dynamic complexes with Ca^2+^-CaM to fulfill their physiological functions^19^.

**Figure 4.**
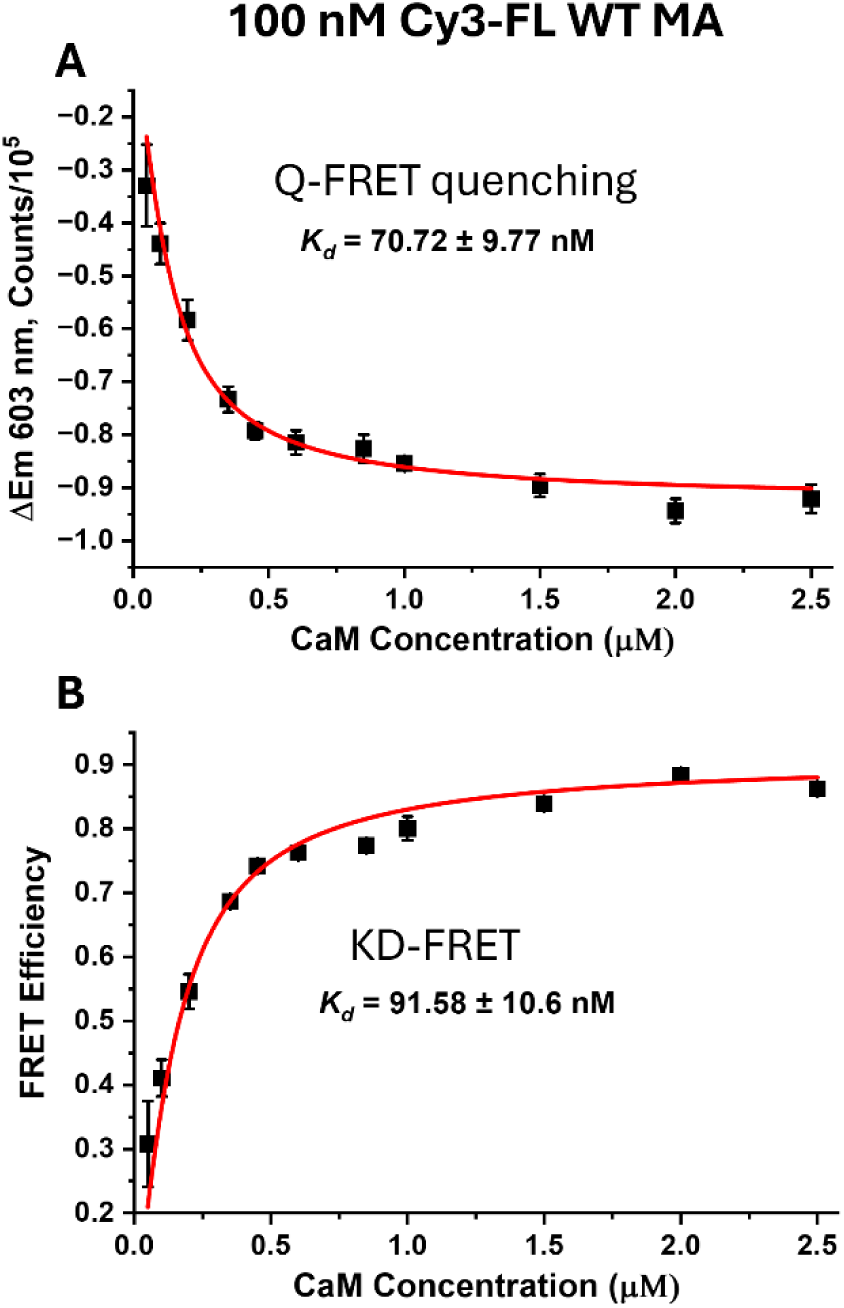
eFRET data analysis for the interaction of 100 nM Cy3-FLWT MA peptide with Cy5-Ca^2+^-CaM at increasing concentrations in the range from 50 nM to 2.5 µM. The background-corrected eFRET data (in Figure 3 A) were used to estimate *K*_d_. The results from fitting the Cy3 donor fluorescence decay at ∼ 603 nm (method of Q-FRET quenching) and fitting the FRET efficiencies (*E*_FRET_) (KD-FRET) to obtain *K*_d_-s are shown in panels (A) and (B), respectively. The estimated *K*_d_-s are shown in the inset. There is a very good agreement between the *K*_d_ values derived using the two methods. All data are averages of triplicate samples—the mean values and standard deviations of three experiments at each condition are shown.

The maximum *E*_FRET_ value estimated using the KD-FRET method for the Cy3-labeled FL WT MA peptide–Cy5-labeled Ca^2+^-CaM complex was subsequently used to calculate the distance between the Cy3 donor and Cy5 acceptor in the MA-Ca^2+^-CaM complex according to the following equation:

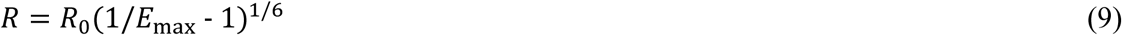

Where, *E*_ma*x*_ is the estimated maximum *E*_FRET_, and *R*_0_ is the Förster distance (radius), which is the specific distance at which the transfer efficiency is exactly 50% (*E* = 0.5) for a given donor-acceptor pair—For the Cy3/Cy5 pair, *R_0_* was set to 53 Å^41, 42^.

Thus, using the estimated *E*_FRET_ of 0.91356 ± 0.01, we calculated a donor–acceptor distance of 35.78 ± 0.76 Å, which is below *R*_0_. This value is consistent with, but also somehow differs from, the NMR structure-based prediction and the AlphaFold3 model of the exact constructs of FL WT MA and Ca^2+^-CaM we used in this study (Supporting Figure S1). The differences may be due to high inherent uncertainty of eFRET distances determination.

#### 3.2.2. The truncated C-terminal region of MA CaM-binding motif (MA-Ct) binds Ca^2+^-CaM with a K_d_ ∼ 65 nM corresponding to a binding energy of about −9.8 kcal/mol

We noticed that the CaM-binding motif of MA contains the sequences AWEKIRLR-----RLKHLVWA (highlighted in cyan in Figure 2), forming the short helices α_1_ and α_2_ of the MA’s CaM-binding motif, which are in contact with the lobes of Ca^2+^-CaM when forming a complex (Figure 1C). Also, a study showed that the tryptophane residues in these regions contribute the binding to Ca^2+^-CaM^43^. The two sequences resemble imperfect repeats, found in other CaM-binding motifs^44^. To learn how the truncation of one of the “repeats” affects the binding to Ca^2+^-CaM, we tested the peptide MA-Ct (Figure 2). Similarly to FL WT MA peptide, we labeled the MA-Ct with Cy3 donor and studied its interaction with Cy5-Ca^2+^-CaM in a titration experiment keeping the Cy3-MA-Ct concentration constant at 100 nM and varying the Cy5-Ca^2+^-CaM concentration in the range of 50 nM to 2.5 µM.

#### 3.2.3. The MA peptide containing the K26N/K27H/K28Q substitutions retains high affinity for Ca^2+^-CaM—K_d_ ∼ 250 nM and ΔG ∼ −9 kcal/mol—indicating that the cluster of positively charged residues is not essential for the formation of the MA–Ca^2+^-CaM complex

Q-FRET quenching and KD-FRET yielded *K*_d_ values of ∼61.92 nM and ∼67.81 nM, respectively (Figure 5). The mean *K*_d_ was therefore ∼65 nM, corresponding to a ΔG of ∼−9.8 kcal/mol. These values are nearly identical to those for the FL WT peptide, suggesting that each half of the MA CaM-binding motif may bind independently. This result contrasts with previous reports that shorter peptides derived from the N- and C-terminal regions of the MA CaM-binding domain (residues 11–29 and 31–46, respectively) bind Ca^2+^-CaM with substantially lower affinity^30^—this was observed for both peptide containing residues 11-29 and 31-46 in the N- and C-terminal regions of MA CaM-binding domain. However, another study reported a *K*_d_ of ∼10 nM for a peptide spanning residues 31–46^45^, which is more consistent with our findings.

**Figure 5.**
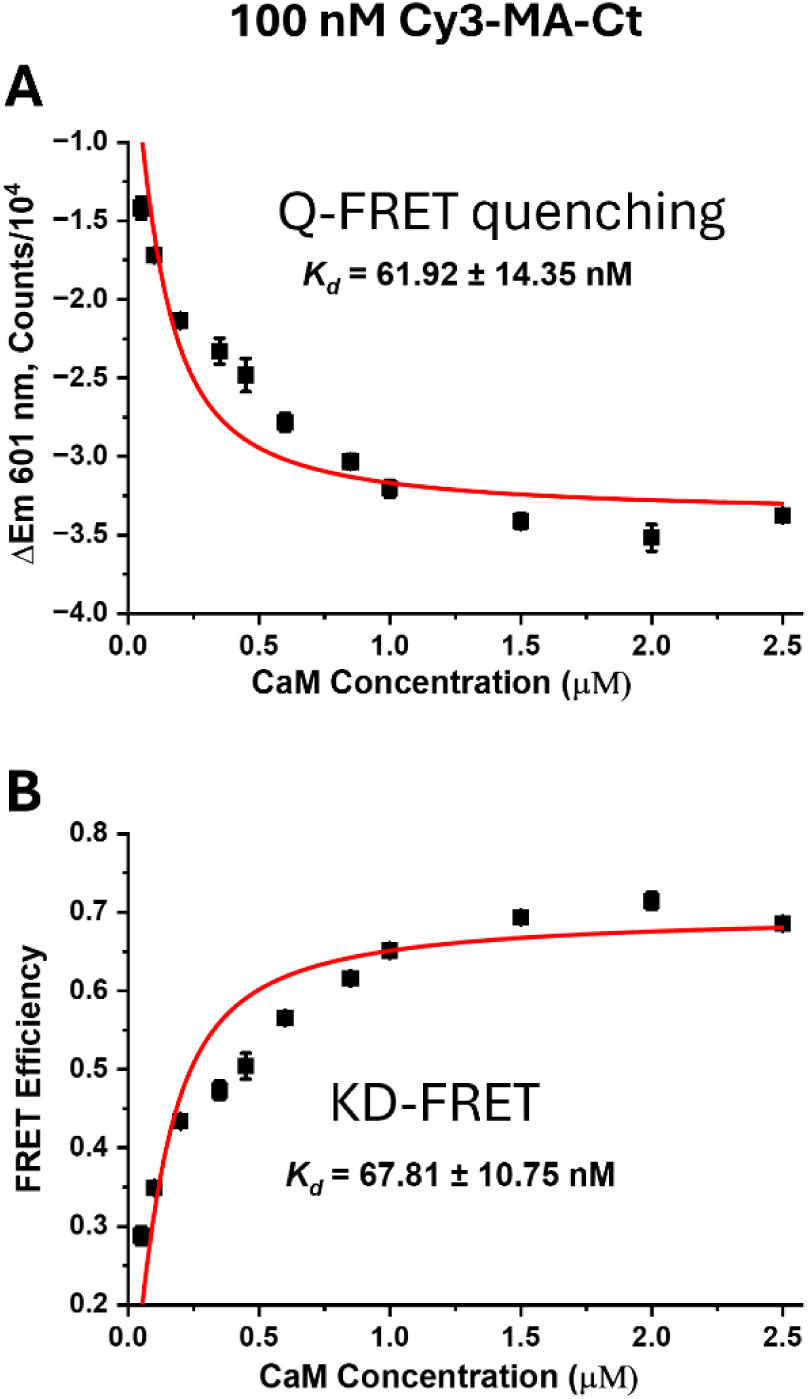
eFRET data analysis for the interaction of 100 nM Cy3-MA-Ct peptide with Cy5-Ca^2+^-CaM at increasing concentrations in the range from 50 nM to 2.5 µM. The background-corrected eFRET data (in Figure 3 B) were used to estimate *K*_d_. The results from fitting the Cy3 donor fluorescence decay at ∼ 601 nm (method of Q-FRET quenching) and fitting the FRET efficiencies (*E*_FRET_) (KD-FRET) to obtain *K*_d_-s are shown in panels (A) and (B), respectively. The estimated *K*_d_-s are shown in the inset. There is a very good agreement between the *K*_d_ values derived using the two methods. All data are averages of triplicate samples—the mean values and standard deviations of three experiments at each condition are shown.

The fit for Cy3-MA-Ct binding to Cy5-Ca^2+^-CaM was poorer than that for Cy3-FL WT MA because of irregular behavior at Cy5-Ca^2+^-CaM concentrations of 200 nM–1 µM (Figure 5). This behavior may reflect MA-Ct binding to a second site in the CaM C-terminal lobe, as previously reported^30^. We are conducting further experiments to clarify this possibility. Consistent with a second binding site, the estimated *E*_max_ of 0.6992 ± 0.01276 yielded a Cy3–Cy5 distance of 46.06 ± 4.7 Å for labels attached to MA-Ct and Ca^2+^-CaM, respectively. This longer distance may represent an average of Cy3-MA-Ct bound to the N- and C-terminal lobes of CaM. Thus, the obtained *K*_d_ could also reflect the binding MA-Ct to the two lobes of CaM. Nevertheless, the binding parameters for MA-Ct strongly suggest that this region of the MA CaM-binding motif forms a strong complex with Ca^2+^-CaM.

Further, although the MA-Ct peptide lacks both the N-terminal region of the MA CaM-binding motif and the positively charged KKK cluster, it bound Ca^2+^-CaM with an affinity comparable to that of FL WT MA, indicating that the KKK cluster is dispensable for this interaction.

Ca^2+^-CaM complex. eFRET titration data collected with 100 nM Cy3-labeled MA-Mut and 50 nM–2.5 µM Cy5-labeled Ca^2+^-CaM (Figure 3C) were analyzed by Q-FRET quenching and KD-FRET (Figure 6A and B). The two methods yielded an average *K*_d_ of ∼250 nM and a calculated *ΔG* of ∼ −9 kcal/mol. Although FL MA-Mut had a higher *K*_d_ than FL WT MA (∼80 nM) when forming a complex with Ca^2+^-CaM, the calculated *ΔG* values differed by less than 1 kcal/mol. Thus, the MA-Mut–Ca^2+^-CaM complex remains stable, indicating that complex formation depends only weakly, if at all, on the HBR K26–K28 lysine cluster. This conclusion is consistent with the strong Ca^2+^-CaM binding of MA-Ct, which also lacks the HBR. To further investigate the slightly reduced affinity of MA-Mut, we used AlphaFold3^36^ to predict the structures of the FL WT MA and FL MA-Mut peptides. The identical predicted structures (Figure 6C) indicate that the substitutions do not cause a major structural perturbation in this MA region, although they may alter local conformational dynamics and modestly reduce binding efficiency.

**Figure 6.**
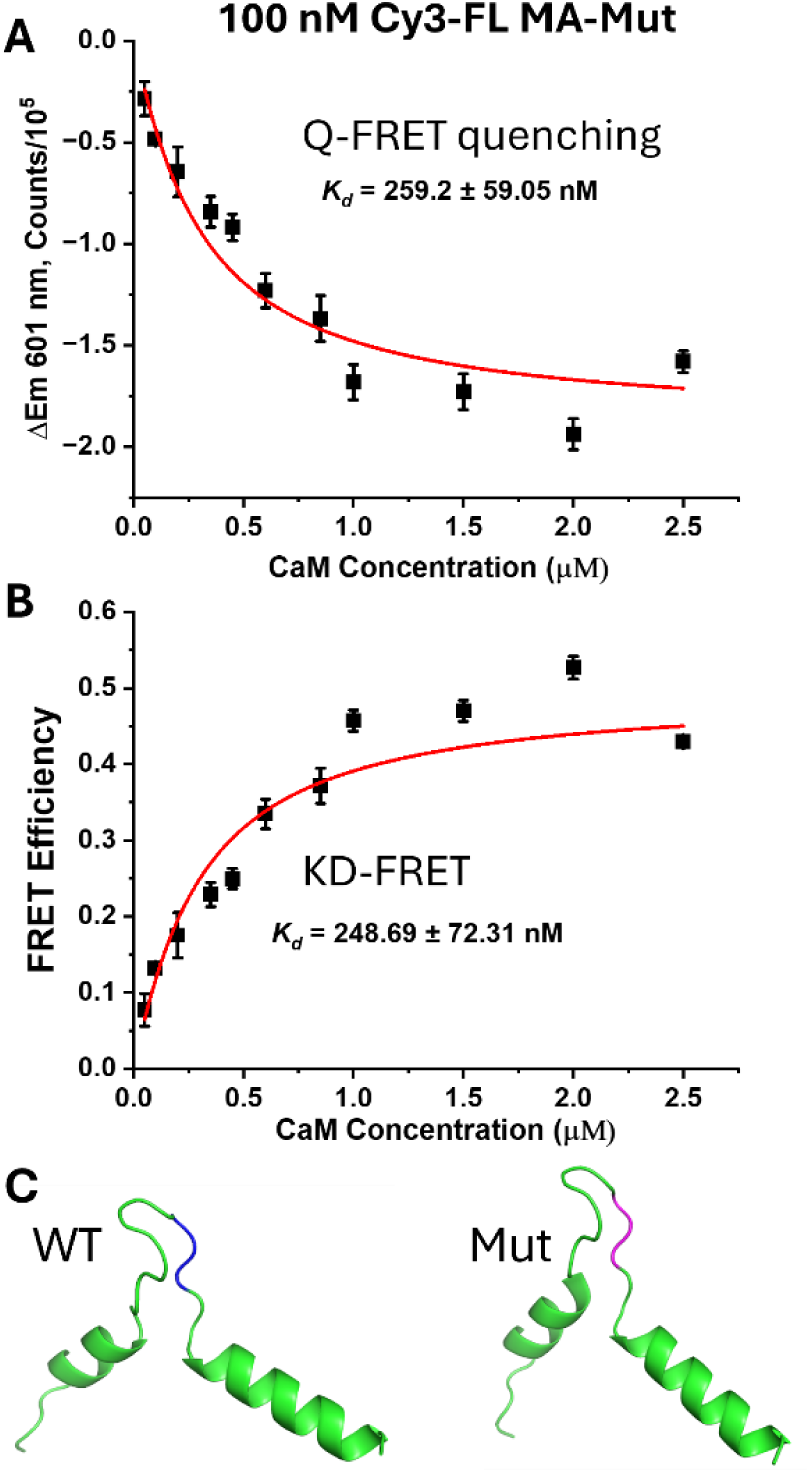
eFRET data analysis for the interaction of 100 nM Cy3-FL MA-Mut peptide with Cy5-Ca^2+^-CaM at increasing concentrations in the range from 50 nM to 2.5 µM. The background-corrected eFRET data (in Figure 3 C) were used to estimate *K*_d_. The results from fitting the Cy3 donor fluorescence decay at ∼ 601 nm (method of Q-FRET quenching) and fitting the FRET efficiencies (*E_FRET_*) (KD-FRET) to obtain *K*_d_-s are shown in panels (A) and (B), respectively. The estimated *K*_d_‘s are shown in the inset. There is a very good agreement between the *K*_d_ values derived using the two methods. All data are averages of triplicate samples—the mean values and standard deviations of three experiments at each condition are shown. The AlphaFold3-predicted structures of the FL WT MA and FL MA-Mut are shown in C—the modeled structures are identical. The region of KKK cluster in WT MA is colored in blue, while the same region containing the K26N/K27H/K28Q substitutions is colored in magenta.

To clarify the contribution of the positively charged HBR residues to Ca^2+^-CaM binding, we examined how the K26N/K27H/K28Q substitutions affect the formation and stability of MA–

Consistent with altered binding, we calculated 53.12±10 Å distance between the Cy3 and Cy5 labels attached to FL MA-Mut and Ca^2+^-CaM, respectively, using the estimated *E*_max_ =0.497±0.028 (KD-FRET). This distance is within the *R_0_* range and is slightly longer than those in the FL WT MA-Ca^2+^-CaM complex, which also supports altered binding conformations.

### 3.3. Reversible MA–Ca^2+^–CaM complex formation is governed by dynamic equilibrium

To determine whether MA–Ca^2+^-CaM complex formation is reversible, we performed competition experiments with two sample series. The first contained 100 nM Cy3-labeled FL WT MA peptide and 100 nM–2.5 µM Cy5-labeled Ca^2+^-CaM. The second had the same composition, but 10 µM unlabeled Ca^2+^-CaM was added after the labeled complex had formed. Unlabeled Ca^2+^-CaM substantially reduced *E*_FRET_ relative to the corresponding samples without competitor (Figure 7, Supporting Figure S2). Consistent with our observations for Vpu–Ca^2+^-CaM^19^, this decrease indicates that unlabeled Ca^2+^-CaM competes for Cy3-MA binding and displaces Cy5-Ca^2+^-CaM from the heterocomplex. These results demonstrate that MA–Ca^2+^-CaM association is reversible and governed by a dynamic equilibrium.

**Figure 7.**
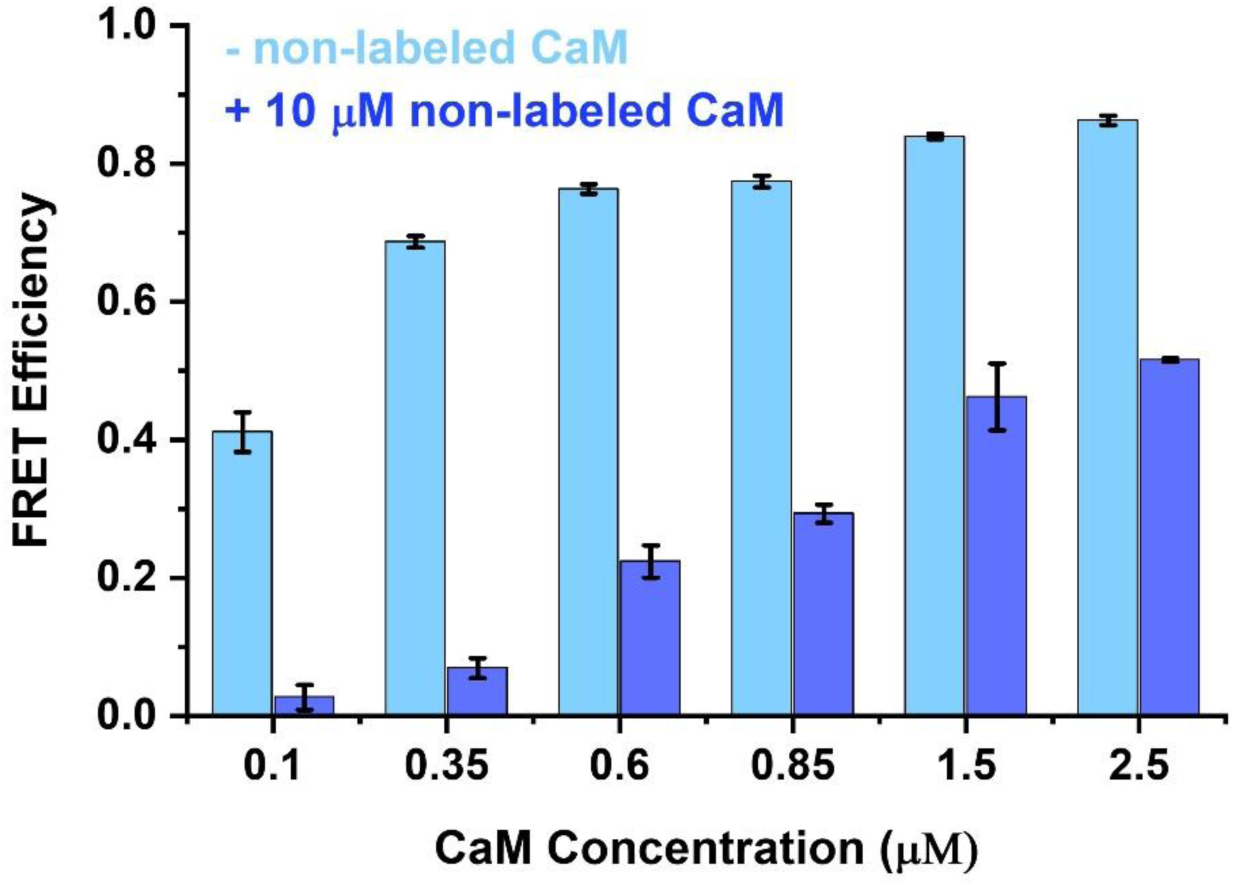
FRET efficiencies for FL WT MA–Ca^2+^-CaM complexes. All samples contained 100 nM Cy3-MA and 100 nM–2.5 µM Cy5-Ca^2+^-CaM. The first series contained only the labeled proteins (light blue bars); the second received 10 µM unlabeled Ca^2+^-CaM after the labeled complex had formed (dark blue bars). Unlabeled Ca^2+^-CaM markedly reduced *E*_FRET_, indicating displacement of Cy5-Ca^2+^-CaM from the Cy3-MA complex. Values are means ± standard deviations from three experiments per condition.

### 3.4. Ca^2+^-CaM adopts a more closed conformation upon binding to FL WT MA peptide

To shed more light on the mechanism of Ca^2+^-CaM-MA complex formation, we conducted DEER study on doubly spin-labeled at residues S39C/A103C Ca^2+^-CaM with and without non-labeled FL WT MA peptide. As expected, just Ca^2+^-CaM was in its open conformation with a long alpha-helix connecting the two globular domains, in agreement with the crystal structure and our earlier DEER measurements^18, 46^. Thus, a well-defined narrow distance distribution with a maximum at about 53 Å between spin-labeled residues S39C/A103C was obtained (Figure 8 A). Next, this distance shifted to shorter values between 18 Å and 30 Å and broader distribution (Figure 8 B), suggesting Ca^2+^-CaM restructured to form the complex with the peptide containing MA’s CaM-binding motif. This is typical restructuring, observed in heterocomplexes of Ca^2+^-CaM with host proteins, as well as in our recent work on the HIV-1 Vpu-Ca^2+^-CaM association^5, 18^. The result is also consistent with the more compact Ca^2+^-CaM conformation when bound to a peptide containing the MA CaM-binding region^12^. The DEER distances between spin-labeled residues S39C and A103C, which we obtained here, are much shorter (by ∼ 10-20 Å) than those predicted based on the NMR structure of ∼ 40 Å (Figure 8 C). However, the DEER distances were more consistent with the AlphaFold3 model, which predicted ∼ 30 Å distance between the C_β_-C_β_ atoms of residues S30C and A102C (Figure 8 D).

**Figure 8.**
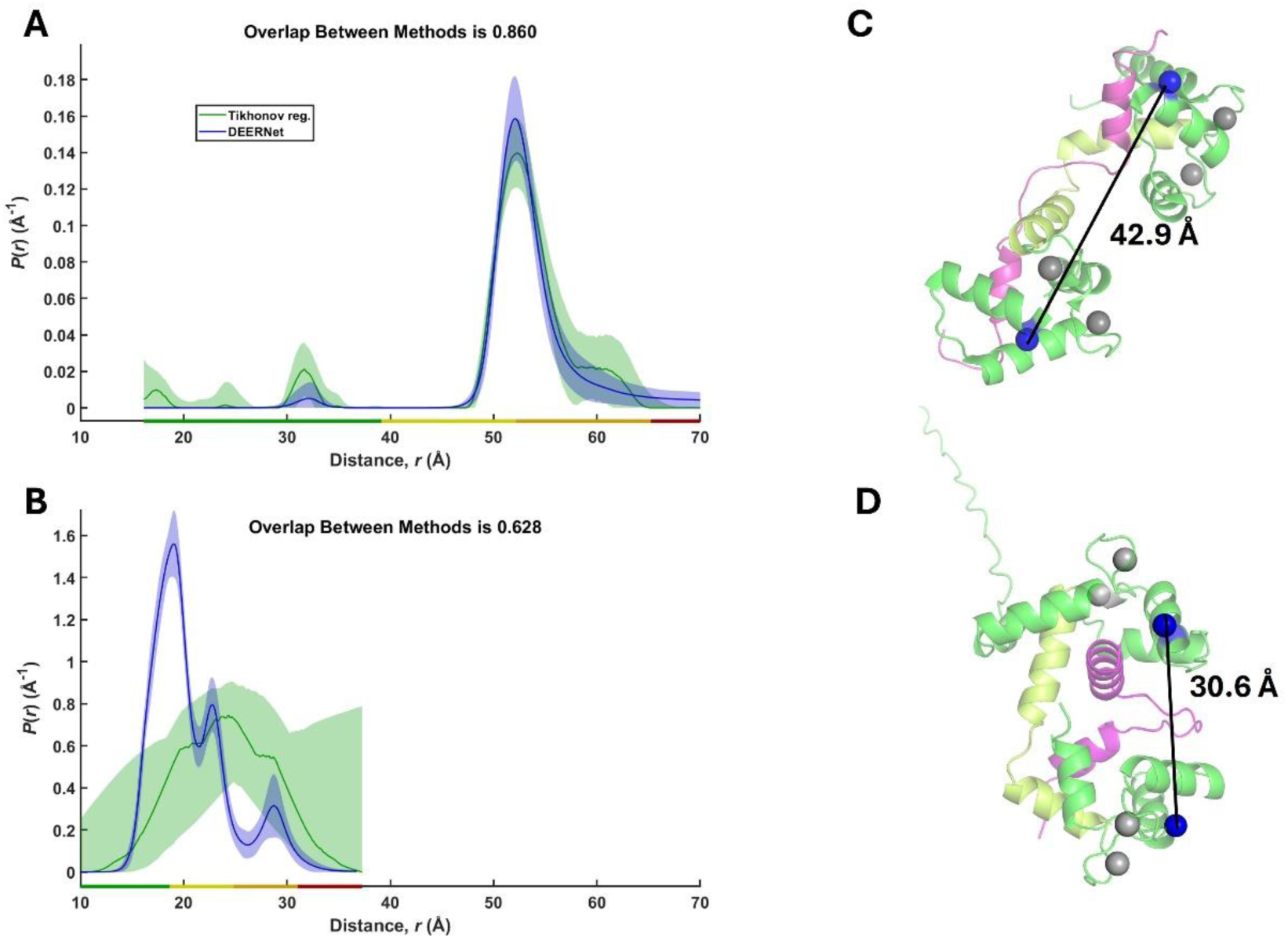
Conformational changes in Ca^2+^-CaM upon binding the MA CaM-binding motif. (A) DEER interspin-distance distribution between spin-labeled residues S39C and A103C in Ca^2+^-CaM without MA peptide. (B) DEER distance distribution for the same residues after binding unlabeled FL WT MA peptide at an approximately 1:1 molar ratio of peptide to Ca^2+^-CaM. Distance distributions reconstructed from the original time-domain DEER signals (Supporting Figure S3) by Tikhonov regularization and DEERNet are in good agreement. MA binding shifts the narrow distribution centered near 53 Å for open Ca^2+^-CaM (A) to substantially shorter distances and broader distributions, consistent with a more closed conformation(B). (C) Predicted distance between the C_β_ atoms (blue spheres) of residues S39 and A103 in the NMR structure of Ca^2+^-CaM bound to the MA CaM-binding motif (PDB 2MGU). Note, there is a shift by one residue in the CaM construct we used (longer by one residue) in this work compared to those used to solve the NMR structure. (D) The AlphaFold3 predicted structure of Ca^2+^-CaM-MA peptide complex. The structure was generated using the Ca^2+^-CaM and FL WT MA peptide from this study. The C_β_ atoms of residues S39C and A102C are blue spheres. The predicted distance of 30.6 Å is shown.

Further, the observed more compact Ca^2+^-CaM conformation in complex with MA compared to its complex with Vpu, as the S39C/A103C distances in the MA-Ca^2+^-CaM complex were shorter by about 10 Å than in the Vpu-Ca^2+^-CaM complex^18^. This could be because the Vpu protein and its C-terminal domain used in our previous study^18^ are simply much larger than the MA peptide used here; therefore one could expect significant steric effects. However, it may well be that the complexes are just different due to unrelated mechanisms of their formation. We also were able to achieve 100% of the Ca^2+^-CaM and MA peptide participation in forming the heterocomplex, judging by the 100% shift to shorter values of the S39C/A103C distances, which was achieved by diluting the system to the low micromolar concentrations where the monomers of MA peptide dominate, and then concentrating the sample to about 50 µm Ca^2+^-CaM and MA peptide concentrations. This is indicative of the 1:1 stoichiometry of the MA-Ca^2+^-CaM complex.

## 4. Discussion

Defining how viral proteins interact with host calmodulin (CaM) is highly relevant to viral calciomics and structural biology^47^. CaM is a ubiquitous, highly conserved eukaryotic Ca^2+^ sensor that regulates hundreds of cellular targets and is universally hijacked during viral infections. Characterizing viral interactions with Ca^2+^-CaM is therefore essential for understanding how viruses manipulate signaling, trafficking, membrane insertion, and other key cellular processes.

CaM is overexpressed in HIV-1-infected cells, and its interactions with HIV-1 proteins are critical to the viral life cycle^8–14^. A detailed understanding of how HIV-1–CaM complexes form could therefore support the development of inhibitors targeting these interactions.

In this study we focused on the interaction of HIV-1 MA protein with Ca^2+^-CaM, as it was proposed that through this interaction the MA domain aids the HIV-1 Gag protein in reaching the plasma membrane at the site of assembly and budding^11^. We used three peptides that correspond to the FL WT CaM-binding motif in MA (Figure 1), as well as a truncated version covering just the C-terminal region of this motif (MA-Ct peptide), and also a modified version of FL CaM-binding motif having the K26K27K28 residues in the HBR within the CaM-binding motif mutated to NHQ amino acids. We wanted to determine the energetics of binding Ca^2+^-CaM to these peptides and elucidate how variations in the MA CaM-binding motif contribute to the heterocomplex formation.

We found that Ca^2+^-CaM forms relatively strong and reversible complex with all the MA peptides studied. Interestingly the Ca^2+^-CaM-FL WT MA complex was characterized with *K*_d_ ∼ 80 nM and *ΔG* ∼ - 9.7 kcal/mols, which are very close to the parameters we found for the HIV-1 Vpu-Ca^2+^-CaM complex^19^. This suggests slightly weaker binding, compared to the complexes of Ca^2+^-CaM with host partners in the range of −10 kcal/mol to −15 kcal/mol, possibly consistent with the need for more dynamic and readily dissociable complexes of MA/Vpu with Ca^2+^-CaM. This is reasonable, given that both MA and Vpu are likely to utilize Ca^2+^-CaM for trafficking to membrane sites; therefore, easy release of Ca^2+^-CaM would facilitate the interaction with lipid bilayer.

The results from our studies agree well with previous binding studies of WT FL MA CaM-binding domain with Ca^2+^-CaM from a different HIV-1 strain, where *K*_d_ of ∼ 25 nM was reported^30^. However, we found that the truncated MA-Ct peptide has affinity for Ca^2+^-CaM, comparable to those of WT FL MA, which agrees with an early study^45^. However, our observations differ from data in a later report, which showed that short peptides corresponding to N- and C-terminal regions of the MA CaM-binding motif have much reduced binding to Ca^2+^-CaM^30^. Thus, these variations could be due to differences in the experimental techniques or MA peptides used.

Because MA is N-terminally myristoylated^27^, previous studies have examined how this posttranslational modification affects MA–CaM interactions. One report suggested that myristoylated MA binds Ca^2+^-CaM with a *K*_d_ of ∼2 µM^11^, corresponding to a relatively weak heteroprotein complex with *ΔG* of ∼ −7.8 kcal/mol. In contrast, our findings clearly demonstrate that MA forms a stable, specific complex with Ca^2+^-CaM and support the earlier proposal^11^ that MA’s myristoylation is not essential for complex formation. However, differences between the binding parameters measured here for non-myristoylated MA and those reported previously for myristoylated MA–Ca^2+^-CaM suggest that this posttranslational modification could affect heterocomplex stability.

One unanswered question in regard to understanding the Ca^2+^-CaM-MA complex formation has been the role of the positive charges in MA HBR, particularly the cluster of positively charged residues central to the CaM-binding motif. Because MA directs Gag to the plasma membrane aiding the assembly of new virions, other important interactions of MA are with membrane lipids, particularly with phosphatidylinositol-(4,5)-bisphosphate [PI(4,5)P2, PIP2], and nucleic acids^48, 49^. The binding to PIP2 and nucleic acids occurs via competitive electrostatic interactions involving the HBR^48, 50^. Yet, it has not been determined whether the MA-CaM and MA-PIP2 association are competitive or complementary, as the MA-PIP2 relies solely on electrostatic attraction, whereas MA-CaM complex was proposed to involve both electrostatic and hydrophobic interactions^11^ and CaM binds also to MA regions outside the stretch of basic 18-33 residues^30^. Our characterization of Ca^2+^-CaM binding to the MA K26N/K27H/K28Q mutant indicates that the native positively charged residues contribute minimally to this protein–protein interaction. This finding suggests that Ca^2+^-CaM and PIP2 or nucleic acids may bind distinct sites on MA rather than compete for the same site. Further competitive binding experiments are needed to address this possibility.

Using DEER distance measurements technique, which is well-suited to study CaM and its complexes^51, 52^, we further captured the changes in Ca^2+^-CaM conformation upon binding to FL WT MA peptide—the distance between spin-labeled residues S39C and A103C in CaM shifted from ca. 5.3 nm to a broader distribution spanning the range from 1.8 nm to 3 nm, confirming the transition from open Ca^2+^-bound conformation–as seen in complexes with human proteins– characterized by the restructuring of the two lobes connected by a long helix into a more compact Ca^2+^-bound state where the binding partner causes the central helix disruption in the middle, exhibiting partial unfolding^5^. Interestingly, the NMR conformation of Ca^2+^-CaM bound to MA CaM-binding motif is less compact^12^ compared to the Ca^2+^-CaM complexes with human proteins^5^. Furthermore, the estimated distances between spin-labeled residues S39C and A103C in the NMR structure with bound MA peptide are significantly longer than our experimental DEER distances.

However, the more compact Ca^2+^-CaM structure produced using AlphaFold3 that more closely matches the complex of Ca^2+^-CaM with host protein agrees with our observations. Thus, it seems that in our study Ca^2+^-CaM adopts more compact conformation to establish contacts with the MA CaM-binding motif.

## 5. Conclusion

Using eFRET and donor-quenching methods, we characterized complex formation between host Ca^2+^-CaM and HIV-1 MA peptides representing the FL WT CaM-binding motif, its truncated C-terminal region, and the FL motif carrying the K26N/K27H/K28Q substitutions. All three constructs bound Ca^2+^-CaM with high affinity (*K*_d_ = 65–250 nM), forming stable and specific yet dynamic and reversible protein heterocomplexes. MA binding also induced substantial restructuring of Ca^2+^-CaM from an extended to a more compact conformation, consistent with observations for Ca^2+^-CaM complexes with other HIV-1 proteins, including Vpu, and with host proteins. These binding parameters help define the properties required of potential inhibitors of the MA–Ca^2+^-CaM complex and advance our understanding of the underlying interaction mechanism.

## Supporting information

Manuscript

## Author Contribution

ASD: experiment, data analysis and interpretation, figures, writing the manuscript; AO: experiment, data analysis and interpretation, figures, writing the manuscript; EH: experiment, data analysis, figures, writing the manuscript; KMM: experiment, data analysis; PPB: experiment, data analysis and interpretation, funds acquisition; ERG: conception, design, experiment, data analysis and interpretation, figures, writing the manuscript, supervision, funds acquisition. All authors contributed to the finalization of the manuscript and approved the final form of the manuscript.

## Acknowledgements

This study was supported by a Gilead Research Scholar in HIV Award to ERG. DEER experiments were conducted at ACERT National Biomedical Resource for Advanced ESR Technology at Cornell University”, Ithaca NY. ACERT is supported by an NIH grant 5R24GM146107.

## Conflict of Interests

The authors declare no conflict of interest.

## Data Availability

Nearly all experimental data produced through this study are shown in the manuscript. The data can be further obtained by contacting the corresponding author upon a reasonable request.

## Notes

### Competing Interest Statement

The authors have declared no competing interest.

