## Supplementary material for "Affinity and Conformations of Ca^2+^-Calmodulin Bound to Peptides Containing the Calmodulin-Binding Motif of HIV-1 Matrix Protein": Manuscript

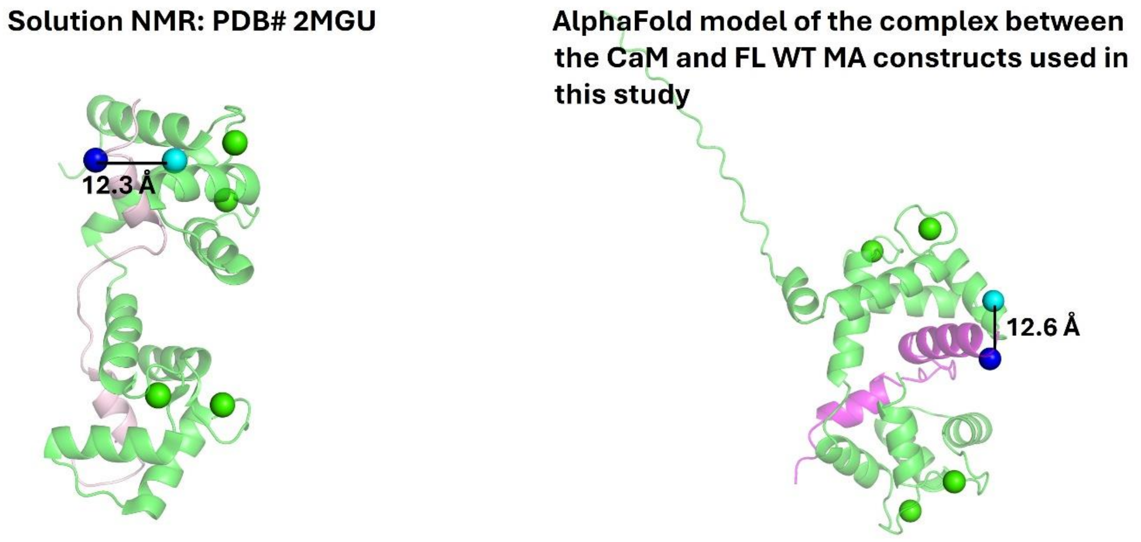


**Figure S1.** Approximate predicted distances between $C_\beta$-$C_\beta$ atoms of the residues in CaM (blue) and MA (cyan), which were labeled with Cy3 and Cy5, respectively. The NMR structure with PDB#2MGU was used. The model structure was produced using AlphaFold3.

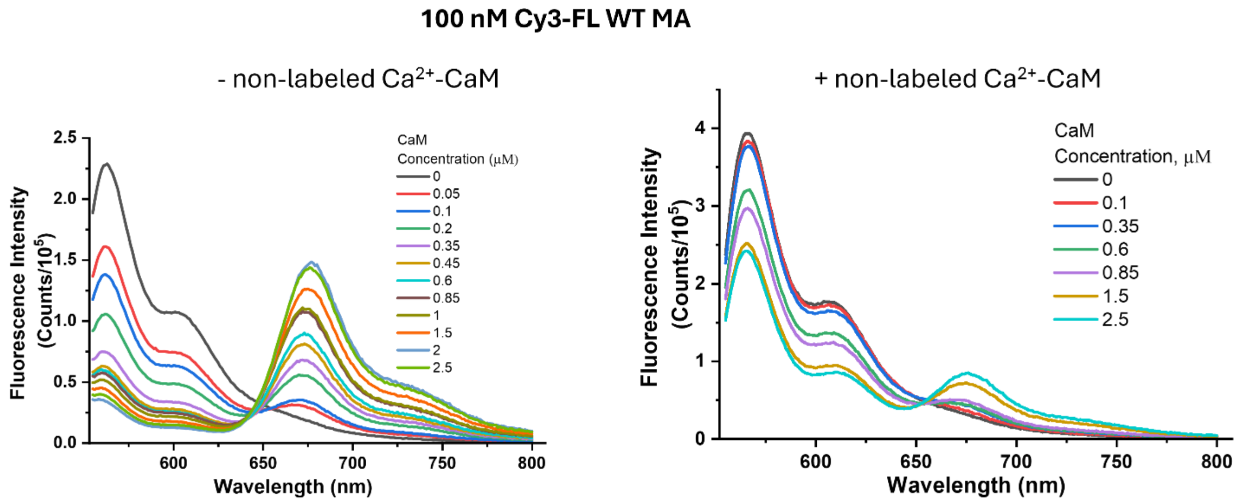


**Figure S2.** eFRET data for the interaction of 100 nM Cy3-FL WT MA with increasing concentrations of Cy5-$Ca^{2+}$-CaM. The Cy5-$Ca^{2+}$-CaM concentrations are shown in the inset, and the data are color-coded. The results without and with 10 μM non-labeled Ca2+-CaM are shown in the left and right panels, respectively. The decrease of FRET signal is apparent in the right panel, compared to the data at the same Cy5-$Ca^{2+}$-CaM concentrations in the left panel.

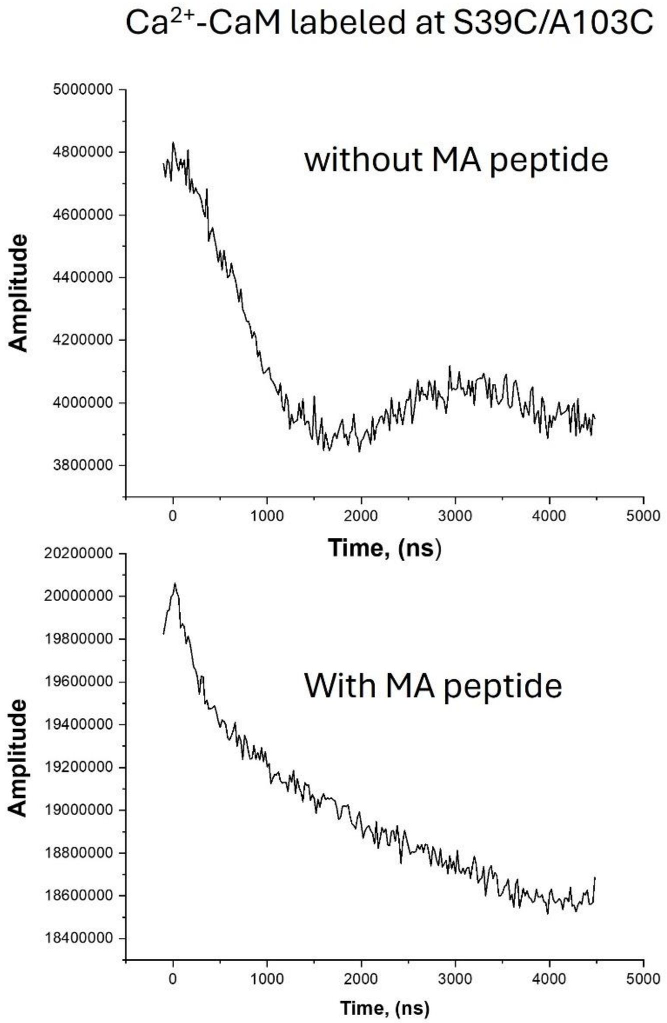


**Figure S3.** Raw DEER data for the samples of spin-labeled $Ca^{2+}$-CaM only (upper panel) and $Ca^{2+}$-CaM bound to FL WT MA peptide.
